# Simulation of zonation-function relationships in the liver using coupled multiscale models: Application to drug-induced liver injury

**DOI:** 10.1101/2024.03.26.586870

**Authors:** Steffen Gerhäusser, Lena Lambers, Luis Mandl, Julian Franquinet, Tim Ricken, Matthias König

## Abstract

Multiscale modeling requires the coupling of models on different scales, often based on different mathematical approaches and developed by different research teams. This poses many challenges, such as defining interfaces for coupling, reproducible exchange of submodels, efficient simulation of the models, or reproducibility of results. Here, we present a multiscale digital twin of the liver that couples a partial differential equation (PDE)-based porous media approach for the hepatic lobule with cellular-scale ordinary differential equation (ODE)-based models. The models based on the theory of porous media describe transport, tissue mechanical properties, and deformations at the lobular scale, while the cellular models describe hepatic metabolism in terms of drug metabolism and damage in terms of necrosis. The resulting multiscale model of the liver was used to simulate perfusion-zonation-function relationships in the liver spanning scales from single cell to the lobulus. The model was applied to study the effects of (i) protein zonation patterns (metabolic zonation) and (ii) drug concentration dependence on spatially heterogeneous liver damage in the form of necrosis. Depending on the zonation pattern, different liver damage patterns could be reproduced, including periportal and pericentral necrosis as seen in drug-induced liver injury (DILI). Increasing the drug concentration led to an increase in the observed damage pattern. A key point for the success was the integration of domain-specific simulators based on standard exchange formats, i.e., libroadrunner for the high-performance simulation of ODE-based systems and FEBio for the simulation of the continuum-biomechanical part. This allows a standardized and reproducible exchange of cellular scale models in the Systems Biology Markup Language (SBML) between research groups.

## 1. Introduction

The liver plays a pivotal role in maintaining metabolic homeostasis. It orchestrates a multitude of metabolic processes that operate on multiple scales and involve complex interactions. Blood is supplied to the liver by two major vessels: the portal vein and the hepatic artery. These vessels branch progressively, culminating in the terminal ends that carry blood to the liver tissues, specifically the hepatic lobules.

Each hepatic lobule, structured in a hexagonal pattern, is a basic functional unit of the liver. Blood inflow occurs at the periphery of the lobule, entering through the portal tracts located at the corners of the lobule. From here, blood flows through the sinusoidal pathways to the central vein located at the core of the lobule. This unique architecture facilitates efficient blood flow and metabolite exchange.

The sinusoids, lined with liver cells called hepatocytes, are the sites of intense metabolic activity. Hepatocytes perform several essential functions, including fat metabolism and detoxification of harmful substances. Their strategic placement along the sinusoids ensures maximum exposure to nutrient-rich blood, allowing for efficient processing and conversion of metabolites.

Hepatic zonation [1], a distinctive spatial organization of metabolic processes within the hepatic lobule, plays a critical role in the liver’s multifaceted functions, including metabolism, detoxification, and bile production. This concept is pivotal in understanding how the liver optimizes its varied tasks across different regions of the lobule.

The lobule can be traditionally segmented into three primary zones, each specializing in specific metabolic activities. Zone 1, or the periportal area, is characterized by high oxygen and nutrient content. This zone is a major hub for oxidative metabolic processes such as gluconeogenesis and fatty acid oxidation. In contrast, Zone 3, or the pericentral area, with its lower oxygen levels, predominantly supports glycolysis, lipogenesis, and notably, the activities of Cytochrome P450 (CYP) enzymes in drug detoxification [2, 3, 4].

CYP enzymes, particularly certain isoforms involved in drug activation and steroid metabolism, exhibit a pronounced zonation pattern within the liver. This distribution is especially notable in the pericentral zone, where these enzymes are highly expressed and selectively induced. This specific localization and induction pattern are crucial for the liver’s capacity to metabolize a wide array of pharmaceuticals and xenobiotics. The selective induction and high expression of CYP isoforms in this zone underscore its vital role in drug metabolism, impacting both drug efficacy and toxicity [5].

This zonal differentiation of CYP activity is not just a structural feature but has significant functional implications. It affects how different drugs are metabolized and detoxified, influencing both therapeutic outcomes and the risk of DILI. Understanding the zonation of CYP enzymes is therefore key to predicting drug behavior within the liver and forms a critical aspect of pharmacological research and liver disease studies.

Mathematical models can be used to describe physiological processes in the liver and to simulate liver diseases. Processes at different scales of the liver can be considered separately. Metabolic models allow to describe the metabolic processes that take place within the liver cells, such as glucose metabolism [6, 7], lipid metabolism [8], central metabolism [9], or drug detoxification [10, 11]. Since liver tissue consists of a fluid saturated porous tissue, porous media approaches can be implemented to simulate tissue behavior [12, 13]. Particular interest has been paid to permeability and microcirculation in porous liver tissue on a lobular scale [14, 15]. The perfusion in liver lobules can therefore be modeled in sinusoids in healthy livers [16, 17] or in fibrotic and cirrhotic livers [18]. An overview for cellular scales is given in [19], for lobular scales in [19, 20, 21].

However, not only must the processes at each scale be considered separately, they also influence each other. Disturbances in perfusion directly affect metabolism in liver cells. Conversely, changes in function, such as the development of liver disease, lead to changes in perfusion [22, 19]. Therefore, multiscale models spanning different size scales of the liver have been presented to simulate the detoxification of substances in the liver at subtissue, tissue, and whole body scales [23, 24]. By combining a porous media approach based on the theory of porous media (TPM) at the lobular scale and systems biology models at the cellular scale, multiscale models are able to describe the function-perfusion relationship with respect to glucose metabolism [25], lipid metabolism [26, 20, 27] or acetaminophen (APAP) detoxification [28, 29].

Most computational models of hepatic drug detoxification focus on APAP which has been modeled by various mathematical approaches [30, 31, 32, 10, 33, 21, 34, 35, 36, 37, 38]. These models can be divided into three approaches: i) Modeling the liver as a single compartment without spatial heterogeneity, either as a model of APAP metabolism alone [10, 35] or coupled to a pharmacokinetic model [30, 32, 36]; ii) Modeling the liver as a linear chain of compartments allowing the study of gradients and zonated metabolism along the sinusoid [31, 23, 33, 34], with most of the models including position-dependent changes in metabolism due to CYP2E1 zonation; iii) Modeling the 2D/3D geometry of the liver lobule explicitly, which allows to study the 2D/3D patterns of zonation and DILI in APAP overdose [37, 38, 28, 29]. Most of these models include position-dependent changes in CYP2E1 and APAP metabolism using different mathematical approaches. Similar 2D/3D approaches have been used to study hepatic drug detoxification by other groups, e.g. [39, 40, 41, 42].

While several 2D/3D models of the liver lobule have been used to investigate the effect of perivenous CYP expression using CYP2E1 as an example, none of the models have systematically investigated the effect of zonation patterns on DILI. Fu et al. studied the effect of periportal, constant, and perivenous zonation patterns on xenobiotic metabolism, but did not study DILI in the form of necrosis [43].

Only the models by Reddyhoff et al. [10] ([BIOMD0000000609](https://identifiers.org/biomodels.db/BIOMD0000000624)) and Sluka et al. [23] (BIOMD0000000624) were easily reusable, with models available in SBML [44].

The aim of this study is to elucidate the histological patterns associated with DILI and hepatotoxicity, particularly in relation to the zonal distribution of metabolic enzymes, with particular emphasis on those involved in drug metabolism. This research focuses on the impact of zonal metabolism, due to the spatial distribution of proteins, on morphological changes in liver tissue using DILI as a model. Despite the critical role of enzymatic zonation in liver function, a comprehensive analysis of its influence on tissue changes remains unexplored. A deeper understanding of this interplay will improve our understanding of the functional changes in hepatic metabolism that result from alterations in enzymatic zonation patterns. This includes understanding how variations in zonation patterns correlate with expected damage manifestations such as necrosis, fibrosis, steatosis, or other pathologies.

## 2. Methods

### 2.1 Model on cellular scale

At the cell scale, an ODE model was used to describe the biochemical SPT reaction network for the irreversible conversion of substrate (S) to product (P) and a toxic by-product (T) via the reaction named S2PT. Substrate and product can be imported into the cell via reversible transporter processes called SIM for S-importer and PEX for P-exporter. T can be detoxified by an irreversible detoxification reaction called TDETOX. All reactions follow Michaelis-Menten kinetics. The model units are time [min], substance [mmol], extent [mmol], and volume [l].

The resulting reaction network and ODE system are shown below:

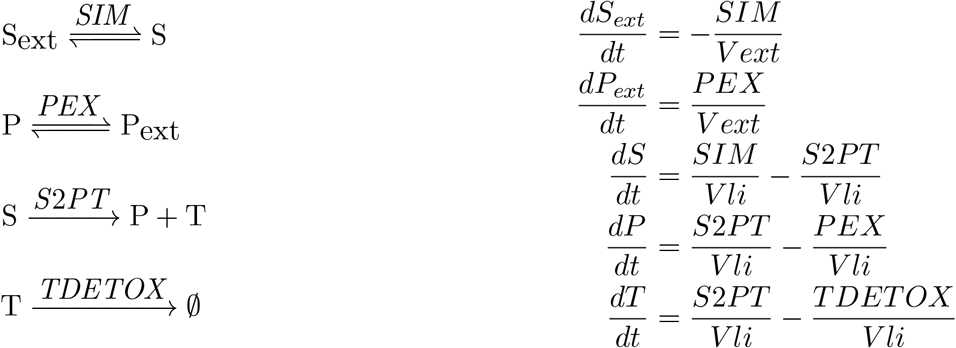

The reaction rates *v* are as follows:

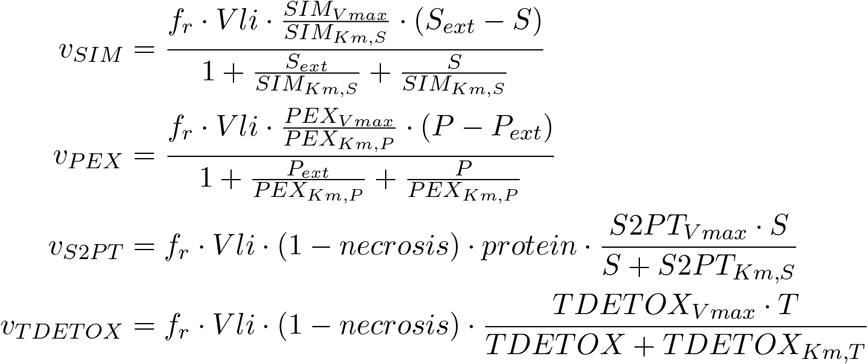

The following additional algebraic rules are defined in the model, defining the relationship between the transport *V max* values and the *V max* value of the *S*2*PT* conversion, the scaling factor *f*_*r*_, and the condition for necrosis (i.e., necrosis occurs if the toxic compound *T* is above *T*_*treshold*_).

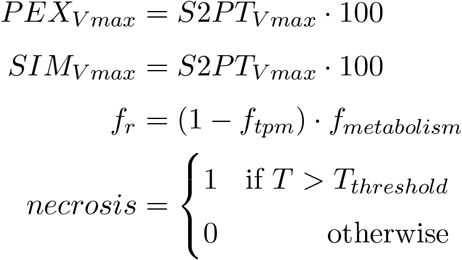

To simulate the influence of different spatial protein zonation patterns on cellular metabolism, six different protein functions were defined depending on the position *p ∈* [0, 1] in the lobulus, where *p* = 0 corresponds to the periportal (PP) and *p* = 1 to the perivenous (PV) position within the lobulus (see Fig. 1).

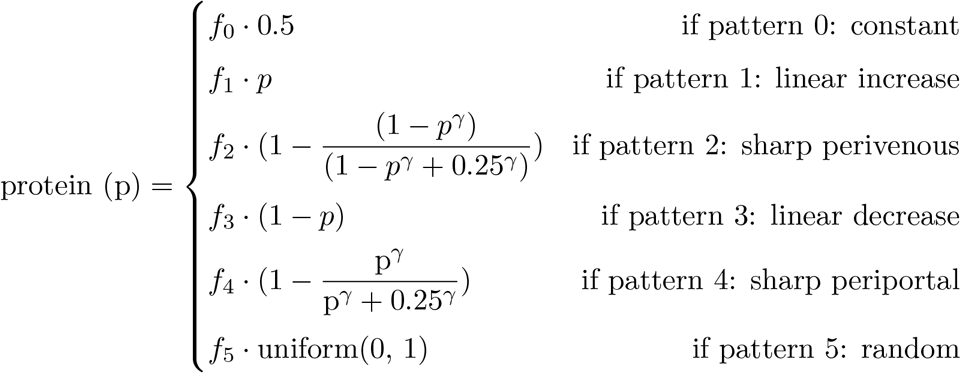

The total amount of protein within the lobules was normalized to the same value for the different patterns using the normalization factors *f*_0_, …, *f*_5_. The function uniform(0, 1) returns a sample from a uniform distribution between 0 and 1. For the random pattern, a seed was used to reproduce the identical random protein pattern for different substrate fluxes. The amount of protein affects the rate of the SPT reaction, i.e., the more protein available at a position *p*, the faster the conversion *S* → *P* + *T* at the given position. The values of all parameters and initial conditions are given in Appendix A.1.

**Figure 1.**
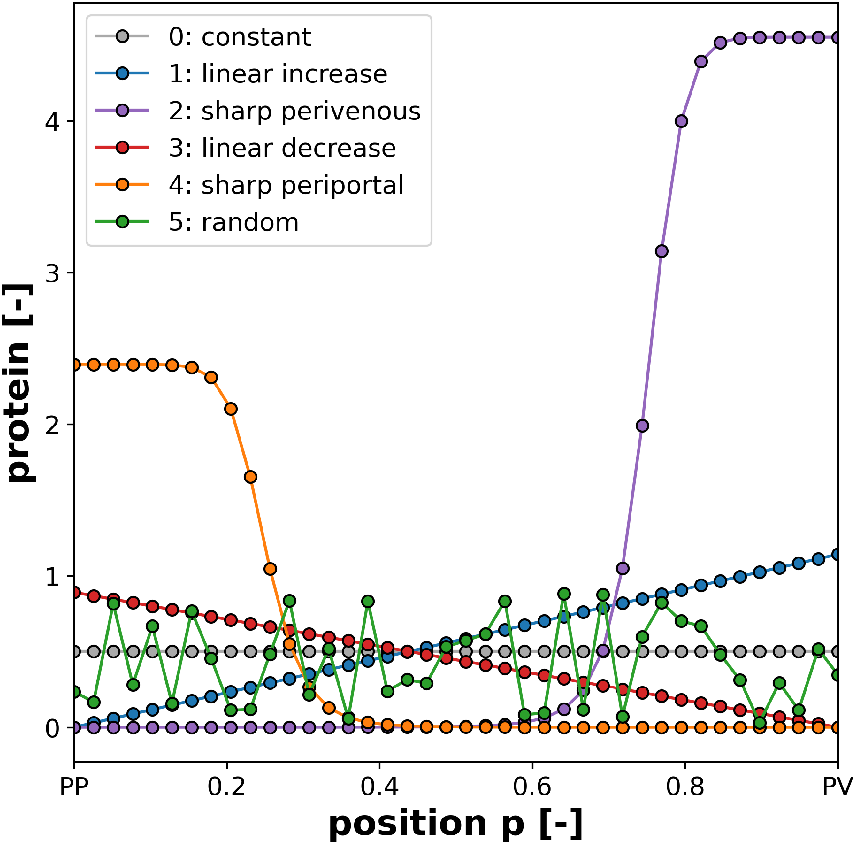
Zonation patterns. Protein amount depending on position *protein*(*p*). Normalization ensures equal protein amount in the lobulus geometry.

The SPT cell model is encoded in the Systems Biology Markup Language (SBML) [45, 46], available under a CC-BY 4.0 license from https://github.com/matthiaskoenig/spt-model, with version 0.5.4 of the model [47] used here. The ODEs are solved numerically with the high performance SBML simulator libroad-runner [48, 49]. For model development and visualization, sbmlutils [50], and cy3sbml [51, 52] were used.

### 2.2. Model on tissue scale

At the tissue level, the liver consists of approximately hexagonal functional units called hepatic lobules. Blood enters the liver lobules at the portal triads at the corners of the lobules and flows to the centrally located central vein. This perfusion through the liver lobules ensures that nutrients, oxygen, and other substances are transported past the liver cells located within the lobules. This is where metabolic processes such as drug metabolism take place. To describe the coupled interaction between structure and fluid, the TPM was used, a homogenization mixture approach based on continuum biomechanics, see [53, 54, 55]. To describe flow and transport of microscopic substances, we extend this approach to include microscopic solutes in blood and liver tissue. To simulate exchange processes between different components, the TPM has been extended with mass exchange processes for miscible [56, 57] and non-miscible substances [58, 59, 60, 61, 62].

In our coupled model we use the TPM for the mathematical calculation of liver tissue containing the immersible phases liver tissue (S) and blood (F) as well as a microscopic substance solved in the blood. This leads to the overall structure of the mixture with all components, where the whole domain *φ* can be described with

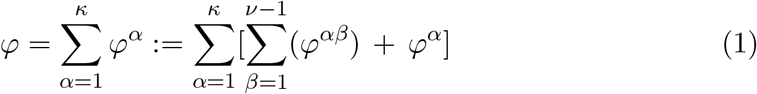

with *κ* = 2 immiscible phases *φ* ^*α*^ = *{*S,F*}* and the microscopic solutes 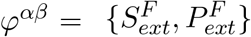.

To describe the physical processes, the momentum balances for each phase and solute can be formulated as

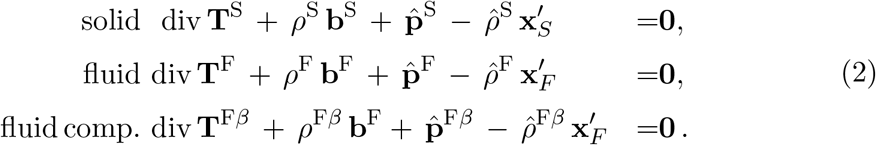

In addition, the mass balances of the individual phases and solutes are added as field equations. Here, exchange terms for the mass exchange between the solutes 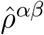 are introduced to describe the metabolic processes.

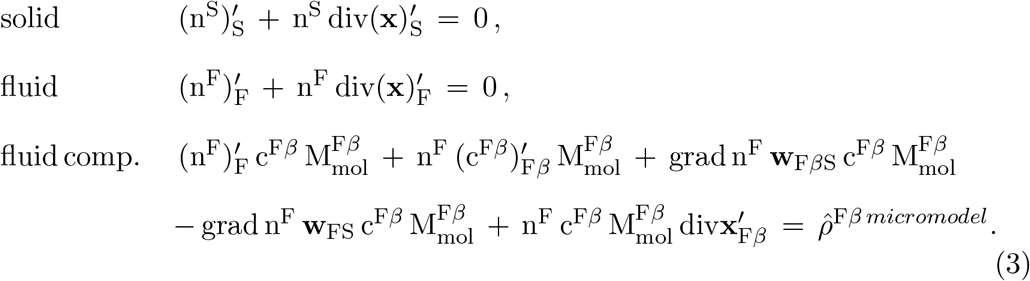

The extended TPM combines the classical mixture theory with the concept of volume fraction, where each phase is defined a volume fraction n^*α*^. The sum of all volume fractions must equal to one, which is denoted in the saturation condition.

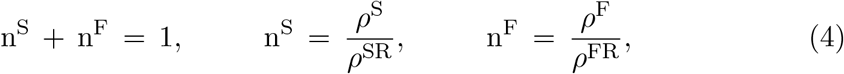

Furthermore, interaction relations between phases have to be considered to incorporate processes in between the separated phases. Therefore, we include relations for the interaction forces **p**^*α*^and mass exchange 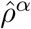 according to the metaphysical principles of Truesdell [63] with

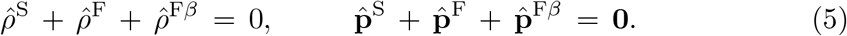

A more detailed derivation of the underlying TPM model can be found in [25, 29]. The continuum-biomechanical model on the lobular scale is solved using the non linear finite element software FEBio [64], which is specifically focused on biomechanics and biophysics. Key structural parameters for the TPM model at the tissue level can be determined from MRI data, cf. [65].

The values of key parameters and initial conditions are given in Appendix A.2.

### 2.3. Multiscale coupling

The coupling of different processes on different scales requires a combination of simulation programs and solvers. For the simulation of the coupled PDE-ODE model, the nonlinear finite element software FEBio [64] was used, embedding the libroadrunner [48, 49] solver for solving the cellular ODEs.

Special attention is given to the correct transfer of variables between scales. The volume V, volume fractions n^*α*^and concentration values c^*αβ*^of the macroscopic domain for a completed time step *t* serve as input to the cellular microsimulations, as shown in Fig. 2C. The microsimulation calculates the source/sink terms as well as their tangents for the time step *t*+*δt* and passes them to the corresponding weak forms of the concentration balance of the macroscopic solver FEBio, where the convergence criterion for the iteration takes place. If convergence is not achieved, the microsimulations are reset and the same time step is attempted with a smaller *δt*. In this context, our approach can be considered as weak or one-way coupling. The existing multiphasic framework of FEBio was extended with the libroad-runner library and invasive coupling code, i.e., getter and setter functions and micromodel initialization. The modified multiphasic model is outsourced from the static FEBio library using the plugin functionality of FEBio [66] to achieve a flexible framework. In this way, the C++ backend of libroadrunner can be compiled as an additional external library without compiling the FEBio core. We follow the object-oriented approach of FEBio by extending the class of material points or integration points by the object of our microsimulation, namely a libroadrunner instance.

**Figure 2.**
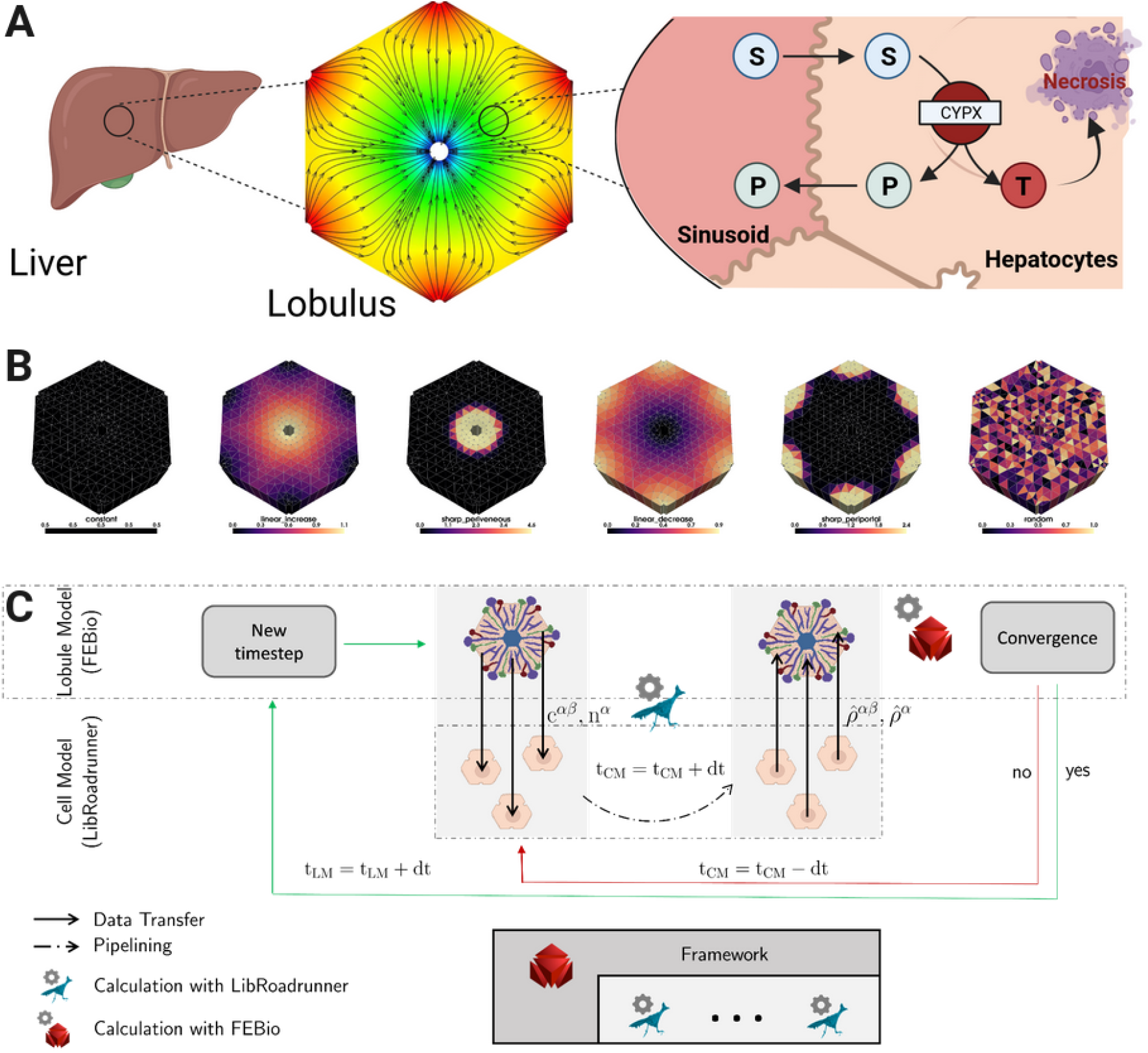
SPT model and model coupling. **A)** Multi-scale model of the liver with organ, lobule and cell scale. The liver is modeled as a set of liver lobules. A single lobule is modeled by an integrated ODE-PDE model. Each point contains a metabolic SPT model that converts S to P with the formation of a toxic by-product. The toxic compound T can cause necrosis if the concentration reaches a threshold value. **B)** Protein zonation pattern of the lobulus: 1. constant amount of protein; 2. linear increase of protein from periportal (PP) to perivenous (PV). 3. strong PV localization of protein. 4. linear decrease of protein from PP to PV. 5. sharp PV pattern. 6. random pattern of protein. **C)** Model coupling and simulation. Outline of the two-scale coupling at the lobular and cellular scale. The values for volume fraction and concentration are calculated on the macro scale and communicated to the micro scale, while source and sink terms are calculated on the micro scale and communicated to the macro scale.

The microsimulations are initialized analogous to the FEBio classes in a global initialization function with the corresponding SPT model for each Gaussian point. Variables like microvolume or volume fraction are used as arguments for the constructor of the microsimulation. The time step of the coupling scheme is shown in Fig. 2C. The time step itself was controlled by the timestepper of FEBio.

A simplified version of the coupling code is provided in Appendix A.4.

### 2.4. Simulation, analysis, visualization

The simulations were performed on the NEC Cluster Vulcan at the HLRS High Performance Computing Center Stuttgart on a Haswell XEON E5-2660v3 with 20 cores and 256 GiB memory.

Visualization and analysis of the FEM simulations were performed in Python using the package porous_media [67], which also contains scripts for pre- and post-processing. Visualizations of the meshes and VTK results were generated using pyvista [68]. Mesh reading and writing was performed using meshio [69].

## 3. Results

### 3.1 Coupled multiscale model

We have developed a multiscale digital twin of the liver that couples a PDE-based porous media approach for the hepatic lobule with cellular-scale ODE-based models for the hepatocytes. An overview of the model and model coupling is provided in Fig. 2. The porous media based models describe transport and tissue mechanical properties and deformations at the lobular scale, while the cellular models describe hepatic drug metabolism and DILI in the form of necrosis (Fig. 2A). The liver is modeled as a set of liver lobules, with a single lobule being modeled as an integrated ODE-PDE model. Each mesh point contains a cellular drug detoxification model that can take up substrate S from the fluid phase and convert S to a product P while forming a toxic by-product T (SPT model). The product P can be exported in the fluid phase, while T can be detoxified via the cells. When the toxic compound T reaches a critical threshold, the corresponding cell can be damaged, resulting in necrosis.

The resulting multiscale model of the liver was applied to simulate perfusion-zonation-function relationships in the liver spanning the scales from the single cell to the lobulus. To illustrate the effect of zonation in drug metabolizing enzymes on potential DILI, the model was simulated with different patterns of protein zonation along the lobule (Fig. 2B).

### 3.2. Cellular simulation

First, the cellular SPT model was studied in isolation to better understand the dependence on substrate concentration S (Fig. 3). Different drug substrate concentrations ranging from 0 to 10 mM were used as initial conditions and the system response was studied over a period of 8 hours. As expected, the concentration of substrate S as well as product P and toxic compound T increases with increasing substrate concentration. At high substrate concentrations (red), the necrosis threshold *T*_*threshold*_ is reached, resulting in cell death. Necrosis is characterized by the release of cell contents into the fluid phase and the inactivation of metabolism. At low substrate concentrations (blue), the liver has sufficient detoxification potential to prevent accumulation of the toxic compound until the threshold for cell damage is reached. DILI can be avoided at low drug substrate concentrations. A clear drug dose dependence can be observed with necrosis occurring above 5 mM and no necrosis below ≤ 5 mM.

**Figure 3.**
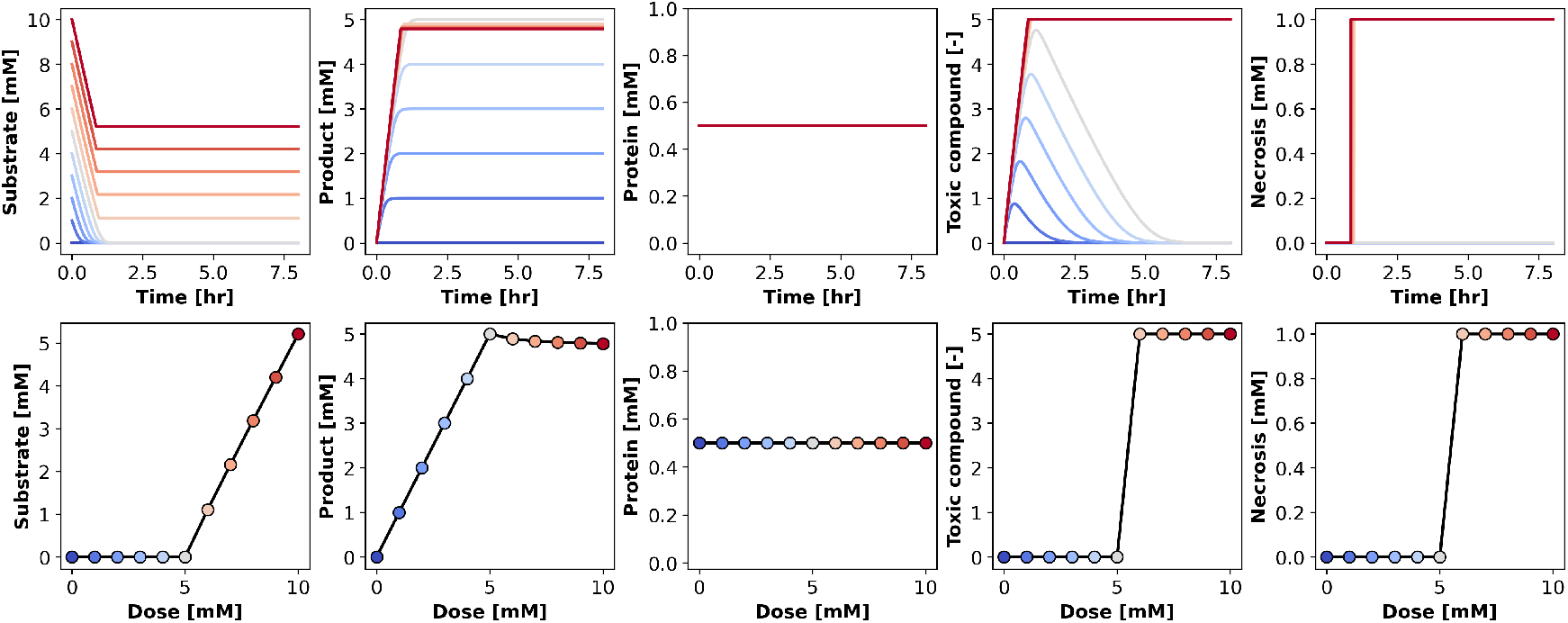
Substrate dependency of SPT metabolism. **A)** Time dependency of drug substrate (S), product (P), protein, toxic compound (T), and necrosis under varying substrate S concentrations. **B)** Substrate dose dependency at steady state.

### 3.3. Multiscale simulation

We then run the coupled multiscale simulations for different zonation patterns and drug (substrate) inputs. Fig. 4 illustrates the behavior of the coupled simulation for the linearly increasing pattern at intermediate substrate flux. Over time, substrate is converted to product with an increase in toxic compound T. After crossing a threshold of 5 mM, local necrosis occurs.

**Figure 4.**
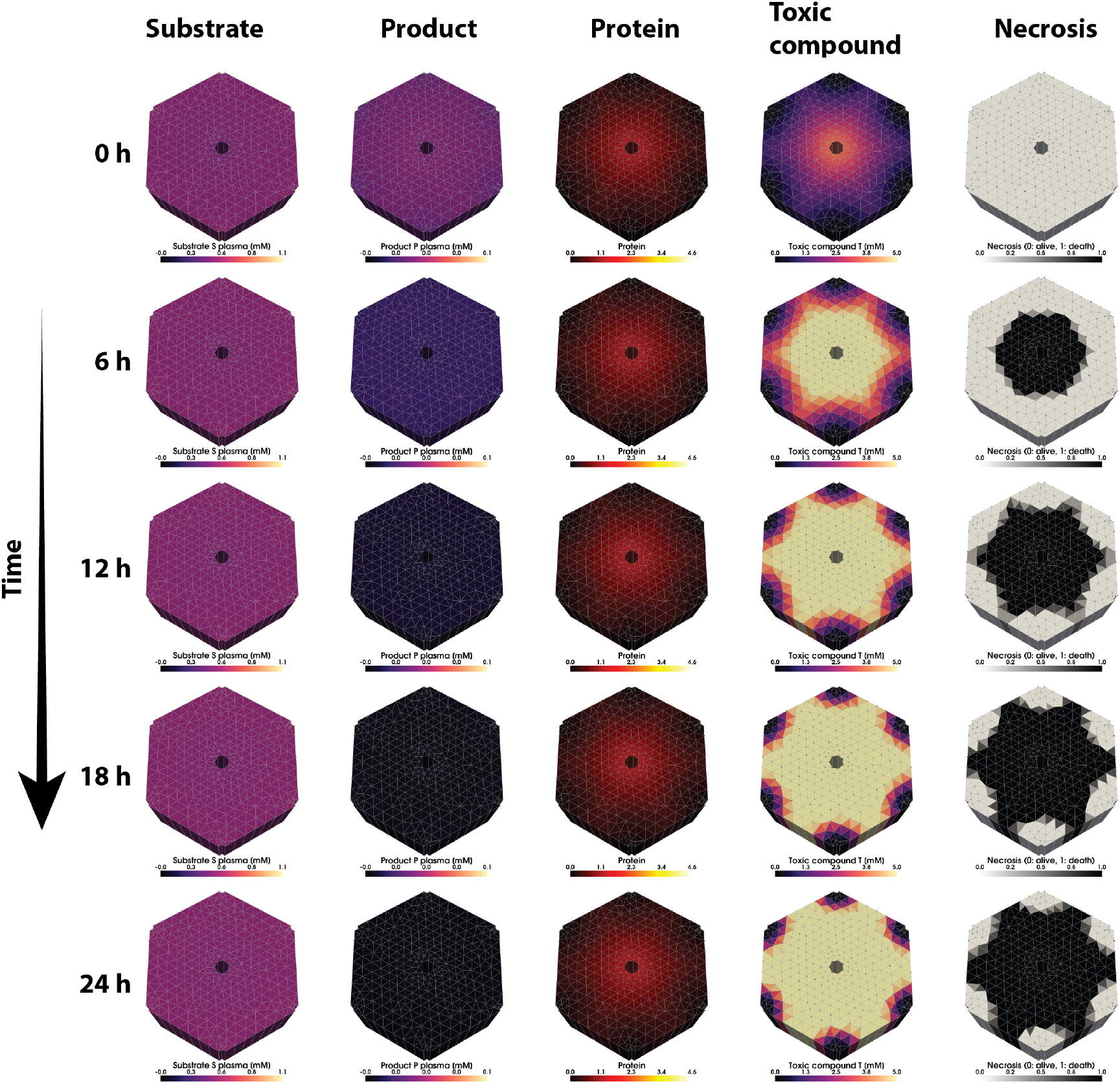
Time course of SPT metabolism (linear increasing pattern and mean substrate flux). Substrate (S), product (P), protein, toxic compound (T) and necrosis over time. Simulation of the PP-PV linearly increasing protein zonation pattern. Simulation for mean substrate flux as boundary condition. T is formed and accumulates over time. Pericentral necrosis forms due to high metabolic rate with high protein concentration (sim014). Simulation for 24 hours. Simulation sim014.

The DILI pattern correlates with the time-independent protein pattern and the concentration of toxic compound T. Over time, the toxic compound accumulates in regions with high protein levels, i.e., pericentrally. The result is pericentral necrosis, with the area affected increasing over time. The concentration profiles of substrate S and product P are different because they are directly coupled to the external concentration at the macro level, where diffusion processes eliminate the gradients. The substrate has a low extraction fraction, with only a small amount cleared in a single passage through the lobule, so the substrate and product gradients are not very strong from periportal to perivenous.

The influence of the six zonation patterns on DILI damage at intermediate drug concentrations is shown in Fig. 5. Six different patterns were investigated: 1. constant amount of protein; 2. linear increase of protein from periportal (PP) to perivenous (PV). 3. strong perivenous localization of protein. 4. linear decrease of protein from PP to PV. 5. strong perivenous pattern. 6. random pattern of protein. The different protein zonation patterns result in very different drug metabolism, toxic compound accumulation, and necrosis patterns. Toxic compound production occurs where protein levels of detoxifying enzymes are high. Over time, necrosis occurs in all patterns, but with very different dynamics. Flat zonation patterns result in a switch-like occurrence of necrosis throughout the lobulus, patterns with predominantly pericentral protein result in pericentral necrosis. Patterns with predominantly periportal protein result in periportal necrosis. The random protein pattern results in random areas of the lobulus being necrotic, corresponding to a high concentration of protein. Over time, the affected area of necrosis increases for periportal, perivenous, and random patterns.

**Figure 5.**
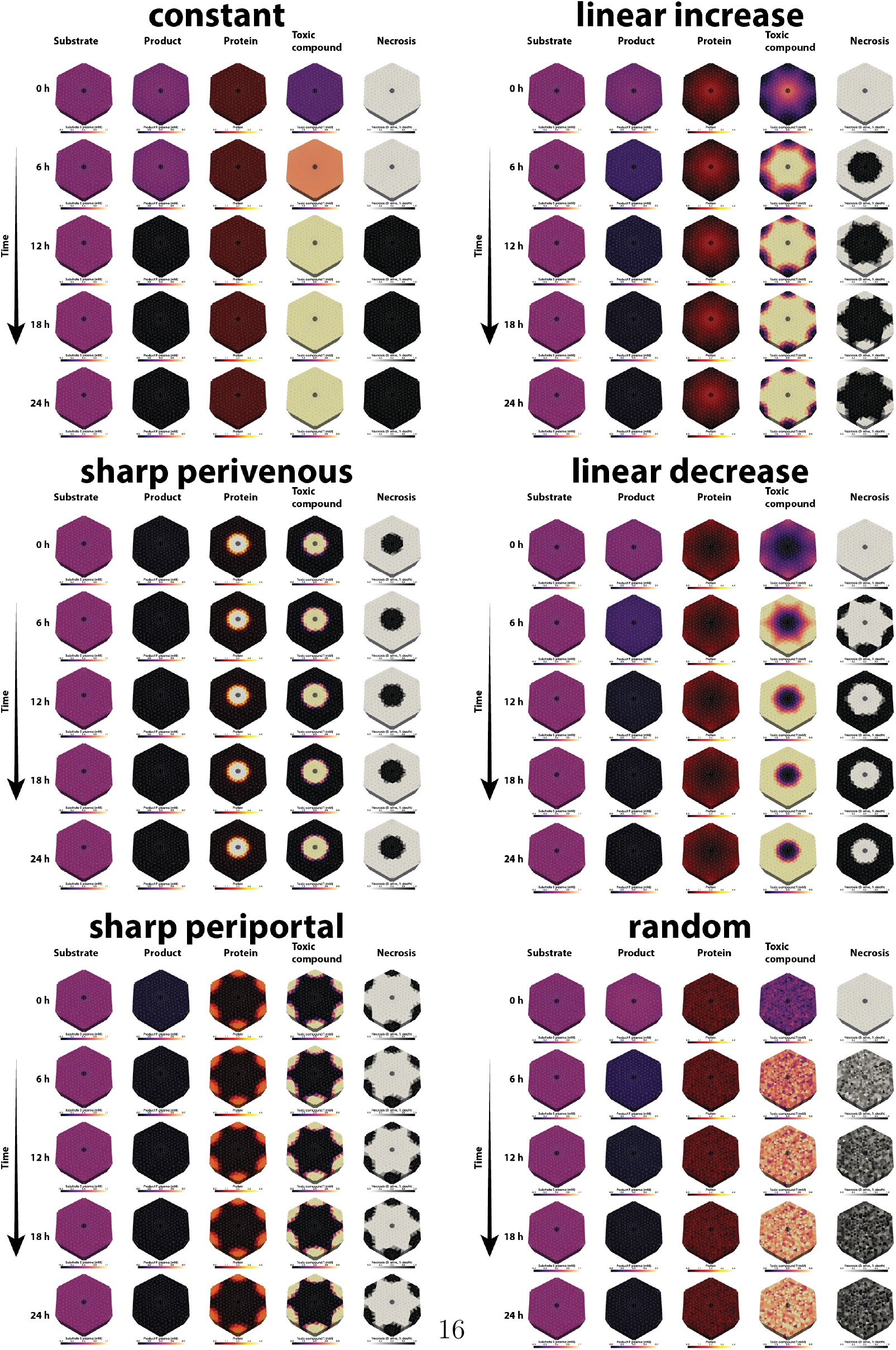
Time course of SPT metabolism depending on zonation patterns. Substrate (S), Product (P), Protein, Toxic Compound (T) and necrosis over time. Simulation of the **A)** constant, **B)** linear increase, **C)** sharp perivenous, **D)** linear decrease, **E)** sharp periportal and **F)** random zonation pattern. Simulation for 24 hours. Simulations sim006, sim014, sim022, sim030, sim038, sim046.

### 3.4. Time dependency of DILI

We then performed a systematic comparison of the effect of protein zonation patterns under different drug (substrate) challenges. Therefore, we first looked at the time evolution of substrate, product, toxic compound, and necrosis area (Fig. 6).

**Figure 6.**
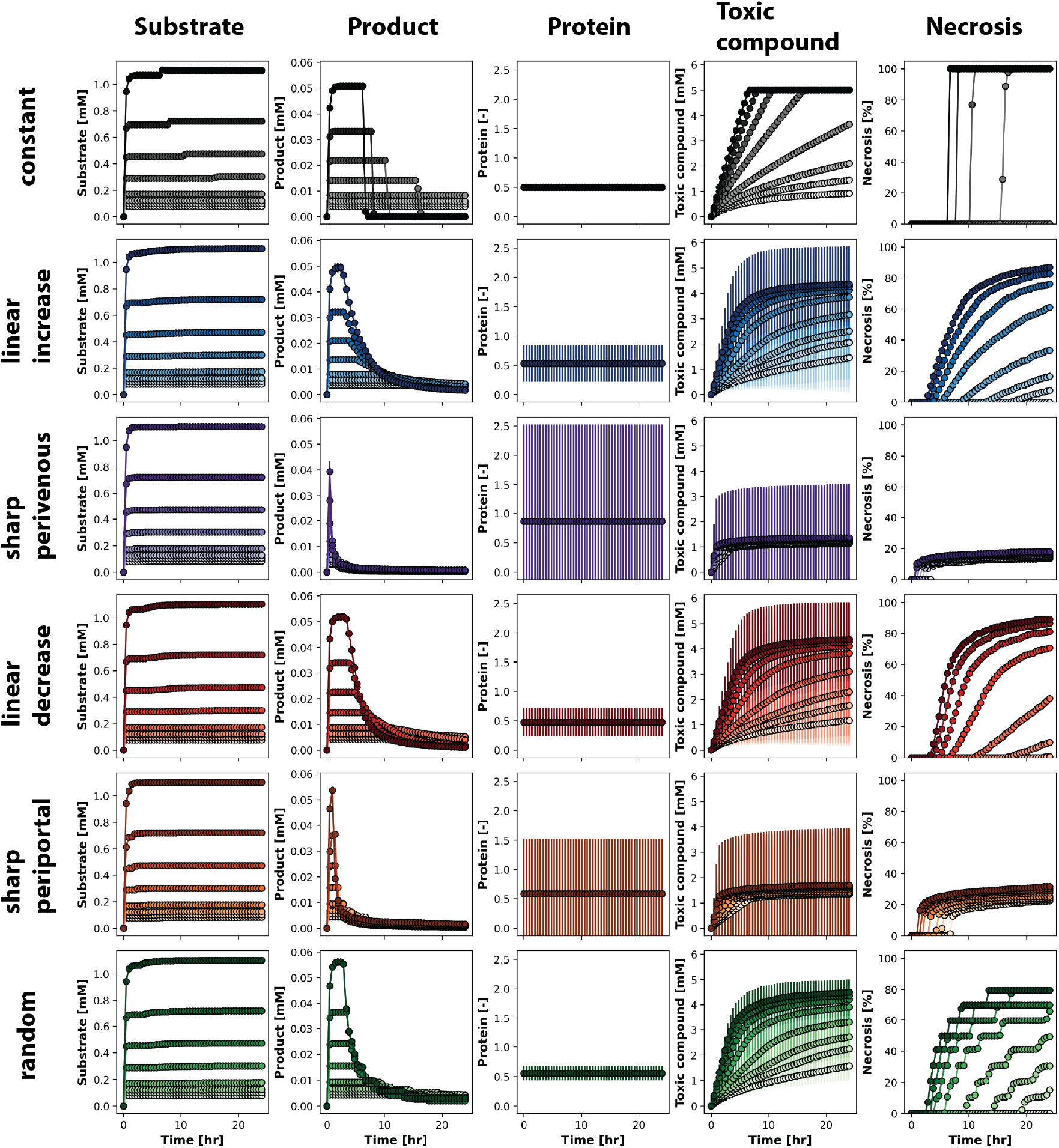
Time dependency of SPT metabolism and necrosis development. Different zonation patterns in rows. Data is mean *±* SD over the lobulus geometry. Necrosis fraction is volume averaged necrosis over the lobulus geometry. Curves correspond to different substrate input fluxes with large substrate fluxes in dark, small substrate fluxes in light colors. Simulation of the **A)** constant (black), **B)** linear increase (blue), **C)** sharp perivenous (purple), **D)** linear decrease (red), **E)** sharp periportal (orange) and **F)** random (green) zonation pattern.

Fig. 6 shows the time dependent accumulation of substrate, product and toxin for different drug concentrations. The values are integrated over the entire lobular geometry. With increasing drug concentration, higher substrate concentrations are reached, resulting in higher product concentrations and faster accumulation of the toxic compound. At higher drug concentrations, necrosis occurs more rapidly and a larger area of liver tissue is affected by DILI, as indicated by the area of necrosis.

As cells die, a small increase in substrate concentration can be observed as substrate is released within the cells. As the area affected by necrosis increases, the product concentration decreases as fewer cells are available for substrate to product conversion. The different zonation patterns show very different dynamics in necrosis. In the case of the constant pattern, there is a rather rapid change from an active to a necrotic state for the entire tissue. For all other protein patterns, even at high drug concentrations, a fraction of the cells survives so that a remaining fraction of the metabolic function can be performed in the lobule. The individual amount of protein for the sharp zonation patterns has a greater variety than for the linear patterns, which is necessary to provide the same average amount of protein across the lobule for the different patterns. The protein remains constant over time as intended. The sharp periportal and perivenous patterns localize most of the protein to a small area affected by necrosis, while the rest of the lobulus is intact. There is not much difference between drug concentrations, but most drug concentrations cause the area of high protein to be necrotic.

The linear and random patterns are much more affected by the different drug concentrations. High drug concentrations result in large areas affected by necrosis, while low concentrations result in smaller areas affected.

### 3.5. DILI patterns

The different protein zonation patterns were challenged with different drug (substrate) fluxes (**Γ**_**S**_) corresponding to increasing drug challenges with the substrate S. In total, 8 different substrate fluxes were tested for the six different protein zonation patterns, resulting in 48 different scenarios (Fig. 7).

**Figure 7.**
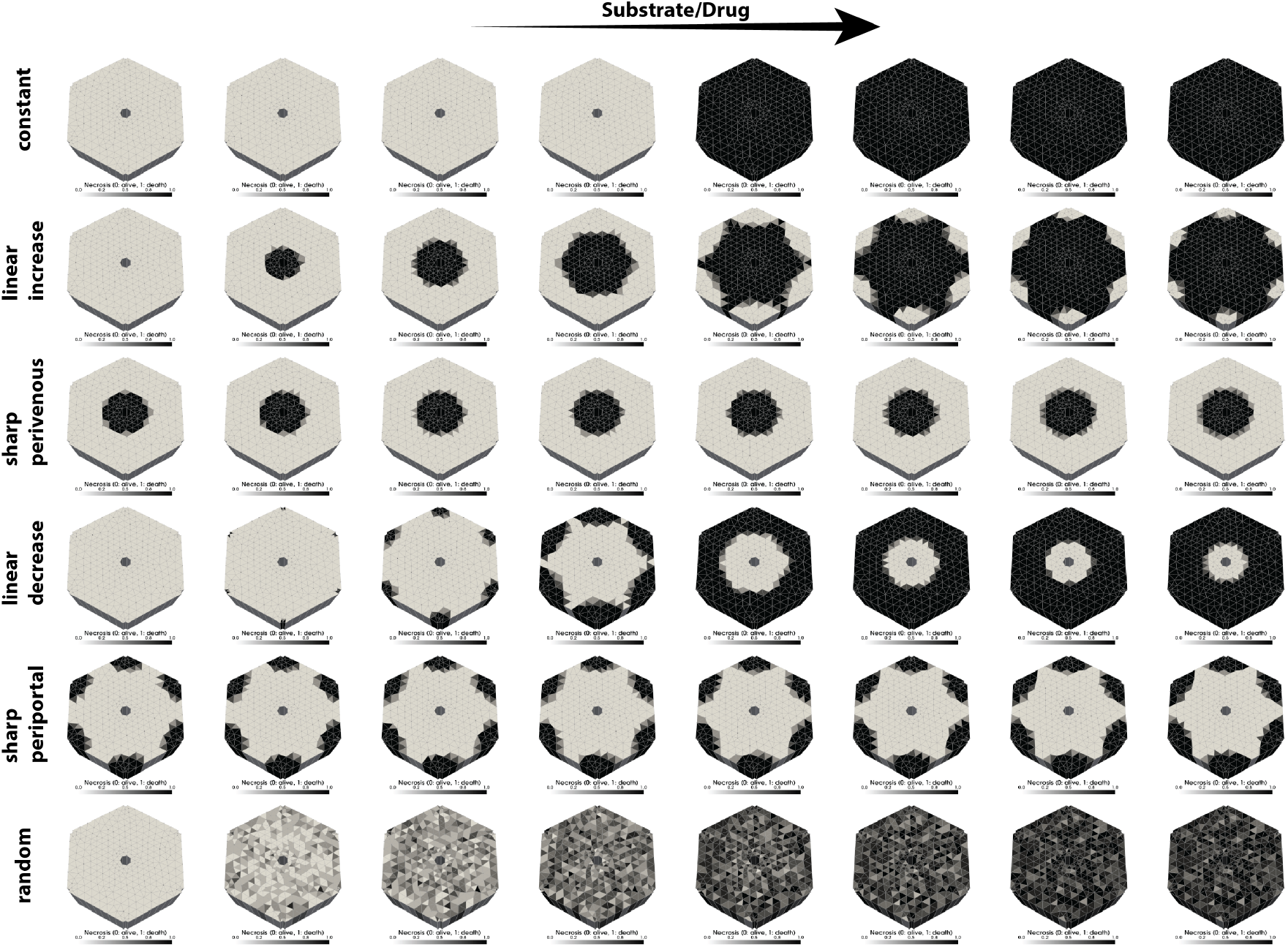
Necrosis pattern depending on zonation pattern and drug (substrate) inflow. Comparison of necrosis patterns at 24 hours for different zonation patterns (rows) and substrate challenges (columns).

The different zonation patterns and different substrate challenges result in highly variable necrosis patterns. As expected, the proportion of necrotic tissue increases as the substrate flow increases from left to right, i.e. the higher the drug dose, the stronger the DILI. For the constant pattern, the absence of intermediate states is striking, so that the metabolic function of the lobule is either completely ongoing or completely terminated across the different substrate fluxes. Periportal protein zonation patterns result in periportal necrosis, pericentral patterns result in pericentral necrosis. The random pattern results in random areas of the lobule affected by necrosis, with the area increasing with increasing drug.

All results are available at https://doi.org/10.5281/zenodo.10933166 [70] with an interactive web application at https://sptmodel.streamlit.app.

## 4. Discussion

In this article, we present a multiscale model of the human liver, including the lobular and cellular scales, based on a PDE-ODE approach. Using the SPT metabolism as a generic example of drug metabolism and DILI, we simulate different zonation patterns to demonstrate their influence on necrosis.

Consistent with other 2D/3D models of APAP detoxification [37, 38], our model predicts pericentrally induced APAP damage (necrosis) as a consequence of pericentral drug-metabolizing enzymes (CYP2E1).

While most DILI is reported as pericentral liver damage because most CYP enzymes are preferentially expressed pericentrally, our model allowed the study of periportal necrosis as a consequence of periportal DILI. Pericentral necrosis due to DILI is the typical phenotype, but cases of periportal necrosis in DILI have been reported. Examples are allyl alcohols [71, 5, 72] or methaprilene [73]. The mainly percicentral DILI damage is consistent with most CYPs being reported to be mainly expressed pericentrally, but evidence exists that some CYP isoforms are mainly located periportal, such as CYP2F2, CYP17A1 or CYP2A12 in mammalian liver [74]. An example of CYP-mediated periportal DILI could be methapyrilene, which causes unusual dose-dependent periportal damage in adult male rats [75, 73, 76, 77], with data suggesting that the toxicity of methapyrilene is predominantly dependent on the CYP2C11 isoform [75].

The concept of dynamic zonation, while not fully integrated in our current micromodel, is a critical aspect to consider for a more comprehensive understanding of liver function. In our study, we simplified the model by assuming spatially variable but temporally constant protein zonation patterns within the hepatic lobule. Although methodologically convenient, this approach overlooks the inherent dynamics of hepatic zonation in response to physiological stimuli. In reality, protein content in hepatocytes is not static; it fluctuates in response to different physiological states, such as feeding or fasting cycles, as well as according to circadian rhythms [78]. These changes in protein expression and distribution can significantly alter hepatic metabolism and detoxification processes. Thus, recognizing and incorporating these dynamic aspects into future models would provide a more accurate and nuanced understanding of liver physiology and its response to different metabolic states. For example, APAP, like many pharmaceutical agents, is thought to exhibit variations in toxicity over a 24-hour (circadian) period [79]. The daily variation in APAP hepatotoxicity (chronotoxicity) may be driven by oscillations in metabolism influenced by the circadian phases of feeding and fasting [80]. In this study, we used constant substrate flows and pressures as boundary conditions for our simulations. However, in a real physiological context, both of these variables are dynamic and influenced by the overall physiology of the body. For example, changes in input substrate concentrations can occur due to factors such as recirculation, distribution within the body, and the involvement of other organs in drug detoxification, particularly renal elimination. Future efforts will aim to incorporate physiologically based pharmacokinetic (PBPK) models [11, 81] to provide dynamic boundary conditions that are more representative of whole-body physiology, as exemplified by the work of Sluka et al. [23].

This study focuses primarily on biomechanical solute transport, incorporating macro-micro coupling for reaction terms. However, it is important to recognize that certain key mechanical processes relevant to liver simulations, such as tissue growth and phase exchange, have not been included. These aspects, although not addressed in this study, are of significant importance and will be the focus of our future investigations. This expansion will improve the comprehensiveness and applicability of our biomechanical models to more accurately understand and simulate liver function.

A significant challenge in current 2D/3D liver modeling approaches lies in the reusability of model components and their adaptability to novel research questions. Our study addresses this issue, with a particular emphasis on enhancing the reproducibility of cellular models. This was achieved by encoding them in SBML [44] and leveraging existing high-performance simulators designed for SBML [48, 49]. Such a methodology not only facilitated the adaptation of our developed model to diverse research inquiries, such as ischemia-reperfusion injury [82], but also provided a streamlined process for updating the cellular models simply by modifying the SBML files. This approach enabled domain experts in tissue-based TPM and cellular ODE models to concentrate effectively on their respective areas of model development.

Future research efforts will incorporate realistic lobular geometries and zonation patterns derived from comprehensive whole slide image analysis [83]. This advancement will allow a more nuanced exploration of how lobular structure influences zonation and drug detoxification processes, thereby deepening our understanding of the structure-function relationship within the liver lobule. In addition, we plan to extend the application of our established workflow for modeling spatial heterogeneity in drug detoxification to specific cases such as APAP. This approach will also be instrumental in investigating the impact of altered metabolic zonation patterns associated with pathologies such as fibrosis [84] and steatosis [85], thereby contributing to a broader understanding of the mechanisms of liver disease.

In conclusion, this study has successfully developed a reusable and expandable liver model specifically designed to study the interplay between perfusion and function and the spatial heterogeneity within the liver lobule. This model represents a significant step forward in understanding liver functionality and its responses to various pharmacological and pathological stimuli.

## Conflict of Interest Statement

The authors declare that the research was conducted in the absence of any commercial or financial relationships that could be construed as a potential conflict of interest.

## Acknowledgements

We thank the FEBio and libroadrunner community, especially Steve Maas, Gerard Ateshian, Lucian Smith, and Herbert Sauro for fruitful discussions and additional help on github and other forums.

## Author contributions

MK and TR designed the study. SG, LL, and MK drafted the manuscript. MK developed the cellular models and generated the SBML. SG, LL, LM, TR, and MK developed the TPM model. SG, LL, LM, JF, and MK designed and implemented the model coupling workflow. SG performed the simulations. MK developed the porous media package, performed the analysis, and prepared the figures. All authors revised the manuscript.

## Funding

SG, TR, and MK were supported by the German Research Foundation (DFG) within the Research Unit Program FOR 5151 “QuaLiPerF (Quantifying Liver Perfusion-Function Relationship in Complex Resection - A Systems Medicine Approach)” by grant number 436883643. LL and TR were supported by Deutsche Forschungsgemeinschaft (DFG, German Research Foundation) under Germany’s Excellence Strategy – EXC 2075 – 390740016. LM, TR, and MK were supported by the DFG by grant number 465194077 (Priority Programme SPP 2311, Subproject SimLivA) and by the Federal Ministry of Education and Research (BMBF, Germany) within ATLAS (grant number 031L0304B). This work was supported by the BMBF-funded de.NBI Cloud within the German Network for Bioinformatics Infrastructure (de.NBI) (031A537B, 031A533A, 031A538A, 031A533B, 031A535A, 031A537C, 031A534A, 031A532B). TR thanks the Deutsche Forschungsgemeinschaft (DFG, German Research Foundation) for support via the project “Hybrid MOR” by grant number 504766766. In addition, TR is supported via the European Union and the German Federal Ministry for Economy and Climate Protection within the framework of the economic stimulus package no. 35c in module b on the basis of a decision by the German Bundestag by the project “DigiTain – Digitalization for Sustainability”. LL and LM are supported by the Add-on Fellowship of the Joachim Herz Foundation.

## Data availability statement

The SPT model generated for this study is available at https://github.com/matthiaskoenig/spt-model. The coupled FEBio-libroadrunner model is available upon reasonable request. All results are available from https://doi.org/10.5281/zenodo.10933166 [70].

## Appendix A. Model details

### Appendix A.1. Parameters of cell model

| parameter | description | value | unit |
| --- | --- | --- | --- |
| $PEX_{Km,P}$ | Km for P export | 0.5 | mM |
| $S2PT_{Km,S}$ | Km for S to P + T conversion | 0.5 | mM |
| $S2PT_{Vmax}$ | Vmax for S to P + T conversion | 0.05 | mmol/l/min |
| $SIM_{Km,S}$ | Km for S import | 0.5 | mM |
| $TDETOX_{Km,T}$ | Km for T detoxification | 0.5 | mM |
| $TDETOX_{Vmax}$ | Vmax for T detoxification | 0.005 | mmol/l/min |
| $T_{threshold}$ | Vmax for T threshold for necrosis | 5.0 | mM |
| $V_{ext}$ | plasma volume | 1.5 | l |
| $V_{li}$ | liver volume | 1.5 | l |
| $f_{metabolism}$ | flag to turn metabolism off | 1.0 | - |
| $f_{tpm}$ | flag for porous media modifications | 0 | - |
| $pattern$ | flag for zonation pattern | $\in (0, 1, 2, 3, 4, 5)$ | - |
| $position$ | norm position between pp and pv | $\in [0, 1]$ | - |
| $protein_{random}$ | random protein value | uniform(0, 1) | - |
| $f_0$ | normalization pattern 0 | 1.0 | - |
| $f_1$ | normalization pattern 1 | 1.1406136 | - |
| $f_2$ | normalization pattern 2 | 4.552802 | - |
| $f_3$ | normalization pattern 3 | 0.8902508 | - |
| $f_4$ | normalization pattern 4 | 2.3939743 | - |
| $f_5$ | normalization pattern 5 | 1.0 | - |
| $\gamma$ | function parameter | 10 | - |
| **state variables** |  |  |  |
| $P[t = 0]$ | product P | 0 | mM |
| $P_{ext}[t = 0]$ | product P (plasma) | 0 | mM |
| $S[t = 0]$ | substrate S | 0 | mM |
| $S_{ext}[t = 0]$ | substrate S (plasma) | 0 | mM |
| $T[t = 0]$ | toxic product T | 0 | mM |

### Appendix A.2. Parameters of tissue model

| **parameter** | **description** | **value** | **unit** |
| --- | --- | --- | --- |
| f_fluid | volume fraction fluid | 0.3 | - |
| f_tissue | volume fraction tissue | 0.7 | - |
| p_in | fluid pressure PP | 1599 | Pa |
| p_out | fluid pressure PV | 933 | Pa |
| perm | constant permeability | 4.25e-11 | m<sup>4</sup>/Ns |
| diffusivity | solute diffusivity | 6.16e-8 | m<sup>2</sup>/s |
| $vS_0$ | substrate flux | -3.797784e-06 | mole/s/m<sup>2</sup> |
| $vS_1$ | substrate flux | -4.6576515e-06 | mole/s/m<sup>2</sup> |
| $vS_2$ | substrate flux | -5.5176906e-06 | mole/s/m<sup>2</sup> |
| $vS_3$ | substrate flux | -7.2377693e-06 | mole/s/m<sup>2</sup> |
| $vS_4$ | substrate flux | -1.0677926e-05 | mole/s/m<sup>2</sup> |
| $vS_5$ | substrate flux | -1.4118084e-05 | mole/s/m<sup>2</sup> |
| $vS_6$ | substrate flux | -1.755824e-05 | mole/s/m<sup>2</sup> |
| $vS_7$ | substrate flux | -2.0938397e-05 | mole/s/m<sup>2</sup> |

### Appendix A.3. Overview simulations

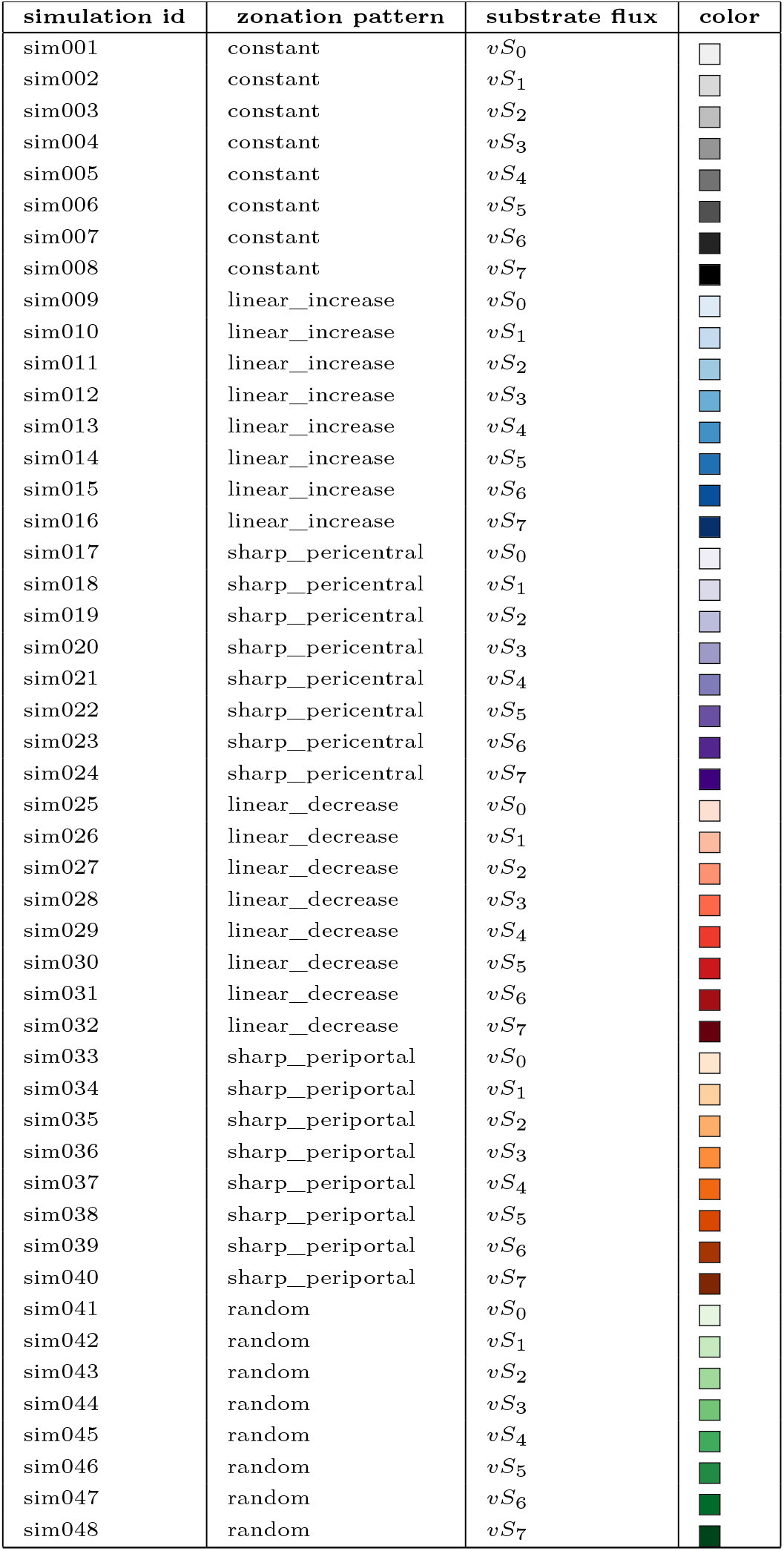

| simulation id | zonation pattern | substrate flux | color |
| --- | --- | --- | --- |
| sim001 | constant | $vS_0$ | |
| sim002 | constant | $vS_1$ | |
| sim003 | constant | $vS_2$ | |
| sim004 | constant | $vS_3$ | |
| sim005 | constant | $vS_4$ | |
| sim006 | constant | $vS_5$ | |
| sim007 | constant | $vS_6$ | |
| sim008 | constant | $vS_7$ | |
| sim009 | linear_increase | $vS_0$ | |
| sim010 | linear_increase | $vS_1$ | |
| sim011 | linear_increase | $vS_2$ | |
| sim012 | linear_increase | $vS_3$ | |
| sim013 | linear_increase | $vS_4$ | |
| sim014 | linear_increase | $vS_5$ | |
| sim015 | linear_increase | $vS_6$ | |
| sim016 | linear_increase | $vS_7$ | |
| sim017 | sharp_pericentral | $vS_0$ | |
| sim018 | sharp_pericentral | $vS_1$ | |
| sim019 | sharp_pericentral | $vS_2$ | |
| sim020 | sharp_pericentral | $vS_3$ | |
| sim021 | sharp_pericentral | $vS_4$ | |
| sim022 | sharp_pericentral | $vS_5$ | |
| sim023 | sharp_pericentral | $vS_6$ | |
| sim024 | sharp_pericentral | $vS_7$ | |
| sim025 | linear_decrease | $vS_0$ | |
| sim026 | linear_decrease | $vS_1$ | |
| sim027 | linear_decrease | $vS_2$ | |
| sim028 | linear_decrease | $vS_3$ | |
| sim029 | linear_decrease | $vS_4$ | |
| sim030 | linear_decrease | $vS_5$ | |
| sim031 | linear_decrease | $vS_6$ | |
| sim032 | linear_decrease | $vS_7$ | |
| sim033 | sharp_periportal | $vS_0$ | |
| sim034 | sharp_periportal | $vS_1$ | |
| sim035 | sharp_periportal | $vS_2$ | |
| sim036 | sharp_periportal | $vS_3$ | |
| sim037 | sharp_periportal | $vS_4$ | |
| sim038 | sharp_periportal | $vS_5$ | |
| sim039 | sharp_periportal | $vS_6$ | |
| sim040 | sharp_periportal | $vS_7$ | |
| sim041 | random | $vS_0$ | |
| sim042 | random | $vS_1$ | |
| sim043 | random | $vS_2$ | |
| sim044 | random | $vS_3$ | |
| sim045 | random | $vS_4$ | |
| sim046 | random | $vS_5$ | |
| sim047 | random | $vS_6$ | |
| sim048 | random | $vS_7$ | |

### Appendix A.4. Pseudocode for coupled simulation

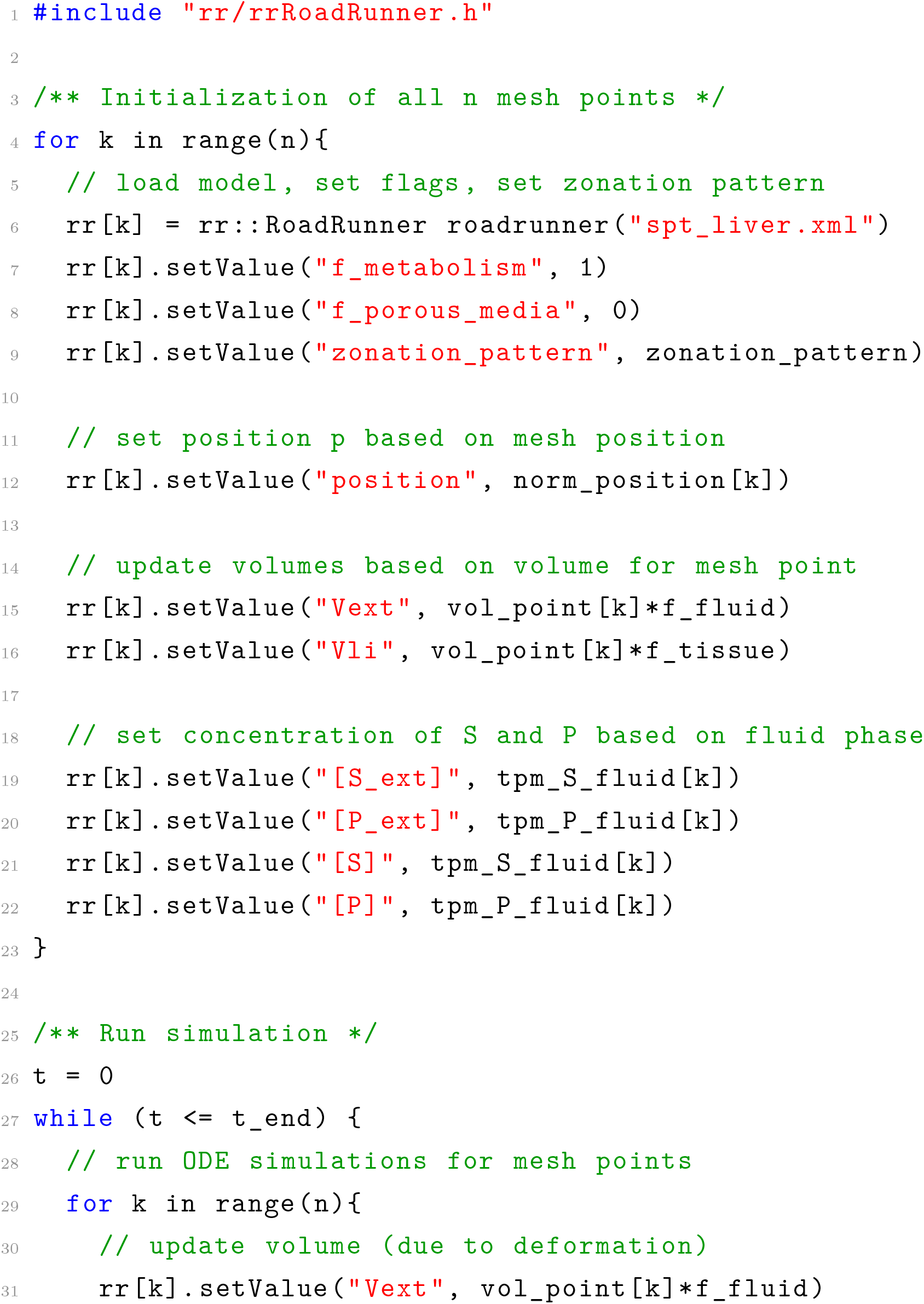
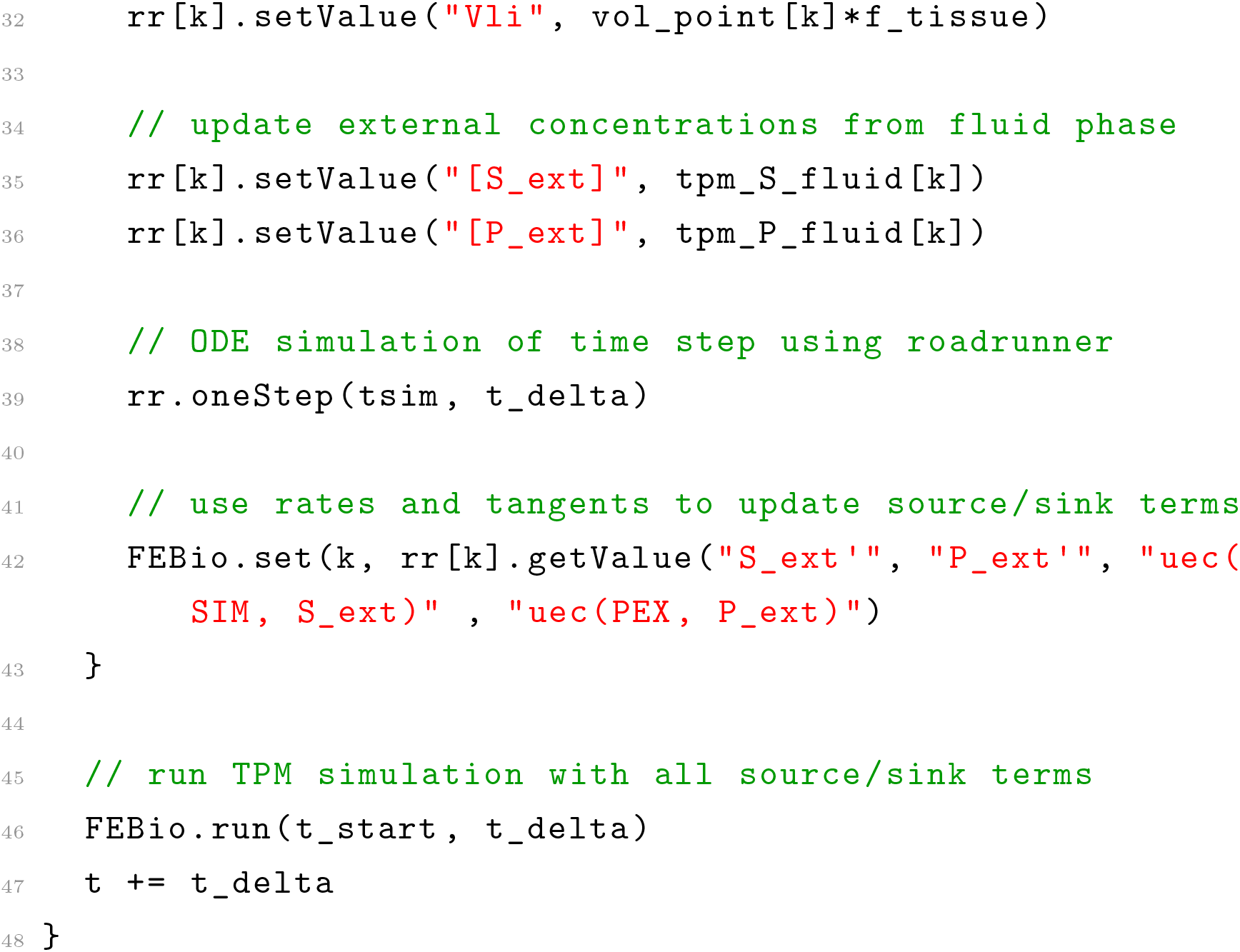

Listing 1: Pseudocode for model coupling. The code illustrates how the cellular SPT models are initialized for every mesh point and the coupled model simulations is run.

## Notes

https://sptmodel.streamlit.app/

### Competing Interest Statement

The authors have declared no competing interest.

### Summary of Updates

All results, images and videos are now available on zenodo https://doi.org/10.5281/zenodo.10933166 In addition, you can navigate through the different zonation patterns and substrate fluxes via the interactive web application at https://sptmodel.streamlit.app. Figures have been updated for clarity. Text has been updated for clarity and feedback from readers has been incorporated.

## References

[1] K. Jungermann, Zonation of metabolism and gene expression in liver, Histochemistry and Cell Biology 103 (2) (1995) 81–91. doi:10.1007/bf01454004.

[2] T. Kietzmann, Metabolic zonation of the liver: The oxygen gradient revisited., Redox biology (2017). doi:10.1016/j.redox.2017.01.012.

[3] S. Ben-Moshe, S. Itzkovitz, Spatial heterogeneity in the mammalian liver, Nature reviews. Gastroenterology & hepatology (2019). doi:10.1038/s41575-019-0134-x.

[4] R. Manco, S. Itzkovitz, Liver zonation, Journal of Hepatology 74 (2) (2021) 466–468. doi:10.1016/j.jhep.2020.09.003.

[5] K. O. Lindros, Zonation of cytochrome P450 expression, drug metabolism and toxicity in liver, General Pharmacology: The Vascular System 28 (2) (1997) 191–196. doi:10.1016/s0306-3623(96)00183-8.

[6] M. König, H.-G. Holzhütter, Kinetic modeling of human hepatic glucose metabolism in type 2 diabetes mellitus predicts higher risk of hypoglycemic events in rigorous insulin therapy, The Journal of biological chemistry 287 (44) (2012) 36978–36989. doi:10.1074/jbc.M112.382069.

[7] M. König, S. Bulik, H.-G. Holzhütter, Quantifying the contribution of the liver to glucose homeostasis: a detailed kinetic model of human hepatic glucose metabolism, PLoS Comput Biol 8 (6) (2012) e1002577. doi:10.1371/journal.pcbi.1002577.

[8] J. Schleicher, R. Guthke, U. Dahmen, O. Dirsch, H. G. Holzhuetter, S. Schuster, A theoretical study of lipid accumulation in the liver-implications for nonalcoholic fatty liver disease, Biochimica et Biophysica Acta (BBA) - Molecular and Cell Biology of Lipids 1841 (1) (2014) 62–69. doi:10.1016/j.bbalip.2013.08.016.

[9] N. Berndt, J. Eckstein, N. Heucke, R. Gajowski, M. Stockmann, D. Meierhofer, H.-G. Holzhütter, Characterization of lipid and lipid droplet metabolism in human hcc, Cells 8 (5) (2019). doi:10.3390/cells8050512.

[10] D. Reddyhoff, J. Ward, D. Williams, S. Regan, S. Webb, Timescale analysis of a mathematical model of acetaminophen metabolism and toxicity, Journal of Theoretical Biology 386 (2015) 132–146. doi:10.1016/j.jtbi.2015.08.021.

[11] A. Köller, J. Grzegorzewski, H.-M. Tautenhahn, M. König, Prediction of survival after partial hepatectomy using a physiologically based pharmacokinetic model of indocyanine green liver function tests, Frontiers in Physiology 12 (2021) 730418. doi:10.3389/fphys.2021.730418.

[12] A. Bonfiglio, K. Leungchavaphongse, R. Repetto, J. H. Siggers, Mathematical modeling of the circulation in the liver lobule, Journal of Biomechanical Engineering 132 (11) (2010) 111011. doi:10.1115/1.4002563.

[13] R. Ahmadi-Badejani, M. Mosharaf-Dehkordi, H. Ahmadikia, An image-based geometric model for numerical simulation of blood perfusion within the liver lobules, Computer Methods in Biomechanics and Biomedical Engineering 23 (13) (2020) 987–1004. doi:10.1080/10255842.2020.1782389.

[14] C. Debbaut, J. Vierendeels, J. H. Siggers, R. Repetto, D. Monbaliu, P. Segers, A 3d porous media liver lobule model: the importance of vascular septa and anisotropic permeability for homogeneous perfusion: the importance of vascular septa and anisotropic permeability for homogeneous perfusion, Comput Methods Biomech Biomed Engin 17 (12) (2014) 1295–1310. doi: 10.1080/10255842.2012.744399.

[15] J. H. Siggers, K. Leungchavaphongse, C. H. Ho, R. Repetto, Mathematical model of blood and interstitial flow and lymph production in the liver, Biomechanics and Modeling in Mechanobiology (2013).

[16] M. Y. Antonov, A. V. Grigorev, A. E. Kolesov, Numerical modeling of fluid flow in liver lobule using double porosity model, in: I. Di-mov, I. Faragó, L. Vulkov (Eds.), Numerical Analysis and Its Applications, Vol. 10187 of Lecture Notes in Computer Science, Springer International Publishing, Cham, 2017, pp. 187–194. doi:10.1007/978-3-319-57099-0\textunderscore18.

[17] F. Bertrand, L. Lambers, T. Ricken, Least squares finite element method for hepatic sinusoidal blood flow, PAMM 20 (1) (2021). doi:10.1002/pamm.202000306.

[18] J. Hu, S. Lü, S. Feng, M. Long, Flow dynamics analyses of pathophysiological liver lobules using porous media theory, Acta Mechanica Sinica 33 (4) (2017) 823–832. doi:10.1007/s10409-017-0674-7.

[19] B. Christ, M. Collatz, U. Dahmen, K.-H. Herrmann, S. Höpfl, M. König, L. Lambers, M. Marz, D. Meyer, N. Radde, J. R. Reichenbach, T. Ricken, H.-M. Tautenhahn, Hepatectomy-induced alterations in hepatic perfusion and function - toward multi-scale computational modeling for a better prediction of post-hepatectomy liver function, Frontiers in Physiology 12 (2021) 733868. doi:10.3389/fphys.2021.733868.

[20] T. Ricken, L. Lambers, On computational approaches of liver lobule function and perfusion simulation, GAMM-Mitteilungen 42 (4) (2019) e201900016. doi:10.1002/gamm.201900016.

[21] H. Ho, S. Means, S. Safaei, P. J. Hunter, In silico modeling for the hepatic circulation and transport: From the liver organ to lobules, WIREs mechanisms of disease 15 (2) (2023) e1586. doi:10.1002/wsbm.1586.

[22] B. Christ, U. Dahmen, K.-H. Herrmann, M. König, J. R. Reichenbach, T. Ricken, J. Schleicher, L. Ole Schwen, S. Vlaic, N. Waschinsky, Computational modeling in liver surgery, Frontiers in Physiology 8 (2017) 906. doi:10.3389/fphys.2017.00906.

[23] J. P. Sluka, X. Fu, M. Swat, J. M. Belmonte, A. Cosmanescu, S. G. Clendenon, J. F. Wambaugh, J. A. Glazier, A liver-centric multiscale modeling framework for xenobiotics, PLoS ONE 11 (9) (2016) e0162428. doi:10.1371/journal.pone.0162428.

[24] J. G. Diaz Ochoa, J. Bucher, A. R. R. Péry, J. M. Zaldivar Comenges, J. Niklas, K. Mauch, A multi-scale modeling framework for individualized, spatiotemporal prediction of drug effects and toxicological risk, Frontiers in pharmacology 3 (2012) 204. doi:10.3389/fphar.2012.00204.

[25] T. Ricken, D. Werner, H. G. Holzhütter, M. König, U. Dahmen, O. Dirsch, Modeling function-perfusion behavior in liver lobules including tissue, blood, glucose, lactate and glycogen by use of a coupled two-scale PDE-ODE approach, Biomech Model Mechanobiol 14 (3) (2015) 515–536. doi:10.1007/s10237-014-0619-z.

[26] T. Ricken, N. Waschinsky, D. Werner, Simulation of steatosis zonation in liver lobule—a continuummechanical bi-scale, tri-phasic, multi-component approach, in: T. Ricken, N. Waschinsky, D. Werner (Eds.), Simulation of Steatosis Zonation in Liver Lobule—A Continuummechanical Bi-Scale, Tri-Phasic, Multi-Component Approach, Vol. 84, Springer International Publishing, 2018, pp. 15–33. doi:10.1007/978-3-319-59548-1\textunderscore2.

[27] L. Lambers, T. Ricken, M. König, Model order reduction (MOR) of function– perfusion–growth simulation in the human fatty liver via artificial neural network (ANN), PAMM 19 (1) (2019) e201900429. doi:10.1002/pamm.201900429.

[28] L. Lambers, N. Waschinsky, T. Ricken, On a multi–scale and multi–phase model of paracetamol–induced hepatotoxicity for human liver, PAMM 18 (1) (2018) e201800454. doi:10.1002/pamm.201800454.

[29] L. Lambers, Multiscale and multiphase modeling and numerical simulation of function-perfusion processes in the liver, dissertation (2023). doi:10.18419/opus-13042.

[30] R. Ben-Shachar, Y. Chen, S. Luo, C. Hartman, M. Reed, H. F. Nijhout, The biochemistry of acetaminophen hepatotoxicity and rescue: a mathematical model, Theoretical Biology and Medical Modelling 9 (1) (Dec. 2012). doi: 10.1186/1742-4682-9-55.

[31] J. G. Diaz Ochoa, J. Bucher, A. R. R. Péry, J. M. Zaldivar Comenges, J. Niklas, K. Mauch, A multi-scale modeling framework for individualized, spatiotemporal prediction of drug effects and toxicological risk, Frontiers in Pharmacology 3 (2013). doi:10.3389/fphar.2012.00204.

[32] E. Leclerc, J. Hamon, I. Claude, R. Jellali, M. Naudot, F. Bois, Investigation of acetaminophen toxicity in hepg2/c3a microscale cultures using a system biology model of glutathione depletion, Cell Biology and Toxicology 31 (3) (2015) 173–185. doi:10.1007/s10565-015-9302-0.

[33] A. K. Smith, B. K. Petersen, G. E. P. Ropella, R. C. Kennedy, N. Kaplowitz, M. Ookhtens, C. A. Hunt, Competing mechanistic hypotheses of acetaminophen-induced hepatotoxicity challenged by virtual experiments, PLOS Computational Biology 12 (12) (2016) e1005253. doi:10.1371/journal.pcbi.1005253.

[34] S. Franiatte, R. Clarke, H. Ho, A computational model for hepatotoxicity by coupling drug transport and acetaminophen metabolism equations, International Journal for Numerical Methods in Biomedical Engineering 35 (9) (Jul. 2019). doi:10.1002/cnm.3234.

[35] S. A. Means, H. Ho, A spatial-temporal model for zonal hepatotoxicity of acetaminophen, Drug Metabolism and Pharmacokinetics 34 (1) (2019) 71–77. doi:10.1016/j.dmpk.2018.09.266.

[36] K. Sridharan, E. A. Ansari, M. Mulubwa, A. P. Raju, A. A. Madhoob, M. A. Jufairi, Z. Hubail, R. A. Marzooq, S. J. R. Hasan, S. Mallaysamy, Population pharmacokinetic-pharmacodynamic modeling of acetaminophen in preterm neonates with hemodynamically significant patent ductus arteriosus, European Journal of Pharmaceutical Sciences 167 (2021) 106023. doi:10.1016/j.ejps.2021.106023.

[37] M. M. Heldring, A. H. Shaw, J. B. Beltman, Unraveling the effect of intra- and intercellular processes on acetaminophen-induced liver injury, npj Systems Biology and Applications 8 (1) (Aug. 2022). doi:10.1038/s41540-022-00238-5.

[38] J. Dichamp, G. Cellière, A. Ghallab, R. Hassan, N. Boissier, U. Hofmann, J. Reinders, S. Sezgin, S. Zühlke, J. G. Hengstler, D. Drasdo, In vitro to in vivo acetaminophen hepatotoxicity extrapolation using classical schemes, pharmacodynamic models and a multiscale spatial-temporal liver twin, Frontiers in Bioengineering and Biotechnology 11 (Feb. 2023). doi:10.3389/fbioe.2023.1049564.

[39] V. Rezania, D. Coombe, J. A. Tuszynski, A physiologically-based flow network model for hepatic drug elimination iii: 2D/3D DLA lobule models, Theoretical Biology and Medical Modelling 13 (1) (Mar. 2016). doi:10.1186/s12976-016-0034-5.

[40] C. A. Hunt, G. E. P. Ropella, L. Yan, D. Y. Hung, M. S. Roberts, Physiologically based synthetic models of hepatic disposition, Journal of Pharmacokinetics and Pharmacodynamics 33 (6) (2006) 737–772. doi:10.1007/s10928-006-9031-3.

[41] L. Yan, G. E. P. Ropella, S. Park, M. S. Roberts, C. A. Hunt, Modeling and simulation of hepatic drug disposition using a physiologically based, multiagent in silico liver, Pharmaceutical Research 25 (5) (2007) 1023–1036. doi: 10.1007/s11095-007-9494-y.

[42] J. Wambaugh, I. Shah, Simulating microdosimetry in a virtual hepatic lobule, PLoS Computational Biology 6 (4) (2010) e1000756. doi:10.1371/journal.pcbi.1000756.

[43] X. Fu, J. P. Sluka, S. G. Clendenon, K. W. Dunn, Z. Wang, J. E. Klaunig, J. A. Glazier, Modeling of xenobiotic transport and metabolism in virtual hepatic lobule models, PLOS ONE 13 (9) (2018) e0198060. doi:10.1371/journal.pone.0198060.

[44] S. M. Keating, D. Waltemath, M. König, F. Zhang, A. Dräger, C. Chaouiya, F. T. Bergmann, A. Finney, C. S. Gillespie, T. Helikar, S. Hoops, R. S. Malik-Sheriff, S. L. Moodie, I. I. Moraru, C. J. Myers, A. Naldi, B. G. Olivier, S. Sahle, J. C. Schaff, L. P. Smith, M. J. Swat, D. Thieffry, L. Watanabe, D. J. Wilkinson, M. L. Blinov, K. Begley, J. R. Faeder, H. F. Gómez, T. M. Hamm, Y. Inagaki, W. Liebermeister, A. L. Lister, D. Lucio, E. Mjolsness, C. J. Proctor, K. Raman, N. Rodriguez, C. A. Shaffer, B. E. Shapiro, J. Stelling, N. Swainston, N. Tanimura, J. Wagner, M. Meier-Schellersheim, H. M. Sauro, B. Palsson, H. Bolouri, H. Kitano, A. Funahashi, H. Hermjakob, J. C. Doyle, M. Hucka, SBML level 3: an extensible format for the exchange and reuse of biological models, Molecular systems biology 16 (8) (2020) e9110. doi: 10.15252/msb.20199110.

[45] M. Hucka, F. T. Bergmann, C. Chaouiya, A. Dräger, S. Hoops, S. M. Keating, M. König, N. L. Novère, C. J. Myers, B. G. Olivier, et al., The Systems Biology Markup Language (SBML): Language Specification for Level 3 Version 2 Core Release 2, Journal of Integrative Bioinformatics 16 (2) (2019) 20190021. doi: 10.1515/jib-2019-0021.

[46] S. M. Keating, D. Waltemath, M. König, F. Zhang, A. Dräger, C. Chaouiya, F. T. Bergmann, A. Finney, C. S. Gillespie, T. Helikar, et al., SBML Level 3: an extensible format for the exchange and reuse of biological models, Molecular Systems Biology 16 (8) (2020) e9110. doi:10.15252/msb.20199110.

[47] M. König, SPT model (Mar. 2024). doi:10.5281/zenodo.10853538.

[48] E. T. Somogyi, J.-M. Bouteiller, J. A. Glazier, M. König, J. K. Medley, M. H. Swat, H. M. Sauro, libRoadRunner: a high performance SBML simulation and analysis library, Bioinformatics (Oxford, England) 31 (20) (2015) 3315–3321. doi:10.1093/bioinformatics/btv363.

[49] C. Welsh, J. Xu, L. Smith, M. König, K. Choi, H. M. Sauro, libRoadRunner 2.0: A High-Performance SBML Simulation and Analysis Library (2022). 2203.01175, doi:10.48550/arXiv.2203.01175.

[50] M. König, sbmlutils: Python utilities for SBML, https://zenodo.org/record/6599299, accessed: 2022-08-23 (Oct. 2021). doi:10.5281/zenodo.5546603.

[51] M. König, A. Dräger, H.-G. Holzhütter, CySBML: a Cytoscape plugin for SBML, Bioinformatics (Oxford, England) 28 (18) (2012) 2402–2403. arXiv: 22772946, doi:10.1093/bioinformatics/bts432.

[52] M. König, N. Rodriguez, matthiaskoenig/cy3sbml: Cy3sbml-v0.3.0 - SBML for Cytoscape, https://zenodo.org/record/3451319, accessed: 2022-08-23 (Sep. 2019). doi:10.5281/zenodo.3451319.

[53] R. Boer, Theory of porous media: highlights in historical development and current state, Springer New York, 2000. doi:10.1115/1.1451169.

[54] R. De Boer, Highlights in the historical development of the porous media theory: toward a consistent macroscopic theory, Applied Mechanics Reviews 49 (4) (1996) 201–262. doi:10.1115/1.1451169.

[55] W. Ehlers, Foundations of multiphasic and porous materials, Springer, 2002. doi:10.1007/978-3-662-04999-0\_1.

[56] W. Ehlers, K. Häberle, Interfacial mass transfer during gas–liquid phase change in deformable porous media with heat transfer, Transport in Porous Media 114 (2) (2016) 525–556. doi:10.1007/s11242-016-0674-2.

[57] T. Ricken, J. Bluhm, Modeling fluid saturated porous media under frost attack, GAMM–Mitteilungen 33 (1) (2010) 40–56. doi:10.1002/gamm.201010004.

[58] M. Robeck, T. Ricken, R. Widmann, A finite element simulation of biological conversion processes in landfills, Waste Management 31 (4) (2011) 663–669. doi:10.1016/j.wasman.2010.08.007.

[59] T. Ricken, A. Sindern, J. Bluhm, R. Widmann, M. Denecke, T. Gehrke, T. C. Schmidt, Concentration driven phase transitions in multiphase porous media with application to methane oxidation in landfill cover layers, ZAMM-Journal of Applied Mathematics and Mechanics/Zeitschrift für Angewandte Mathematik und Mechanik 94 (7-8) (2014) 609–622. doi:10.1002/zamm.201200198.

[60] T. Ricken, A. Thom, T. Gehrke, M. Denecke, R. Widmann, M. Schulte, T. C. Schmidt, Biological driven phase transitions in fully or partly saturated porous media: A multi-component FEM simulation based on the theory of porous media, in: Views on Microstructures in Granular Materials, Springer International Publishing, 2020, p. 157–183. doi:10.1007/978-3-030-49267-0_8.

[61] S. M. Seyedpour, A. Thom, T. Ricken, Simulation of contaminant transport through the vadose zone: A continuum mechanical approach within the framework of the extended theory of porous media (etpm), Water 15 (2) (2023) 343. doi:10.3390/w15020343.

[62] S. M. Seyedpour, C. Henning, P. Kirmizakis, S. Herbrandt, K. Ickstadt, R. Doherty, T. Ricken, Uncertainty with varying subsurface permeabilities reduced using coupled random field and extended theory of porous media contaminant transport models, Water 15 (1) (2023) 159. doi:10.3390/w15010159.

[63] C. Truesdell, Rational Thermodynamics, Springer New York, 1984. doi:10.1007/978-1-4612-5206-1.

[64] S. Maas, B. Ellis, A. Ga, W. JA, FEBio: finite elements for biomechanics., J Biomech Eng. 134 (1) (2012) 011005. doi:10.1115/1.4005694.

[65] S. M. Seyedpour, M. Nabati, L. Lambers, S. Nafisi, H.-M. Tautenhahn, I. Sack, J. R. Reichenbach, T. Ricken, Application of magnetic resonance imaging in liver biomechanics: A systematic review, Frontiers in Physiology 12 (2021) 733393. doi:10.3389/fphys.2021.733393.

[66] S. Maas, S. LaBelle, A. GA, W. JA, A plugin framework for extending the simulation capabilities of FEBio., Biophys J. 115 (9) (2018) 1630–1637. doi: 10.1016/j.bpj.2018.09.016.

[67] M. König, porous_media: python utilities for porous media analysis and visualization (Feb. 2024). doi:10.5281/zenodo.10607553. URL 10.5281/zenodo.10607553

[68] C. B. Sullivan, A. A. Kaszynski, Pyvista: 3D plotting and mesh analysis through a streamlined interface for the visualization toolkit (VTK), Journal of Open Source Software 4 (37) (2019) 1450. doi:10.21105/joss.01450.

[69] N. Schlömer, meshio: Tools for mesh files (Jan. 2024). doi:10.5281/zenodo.1288334. URL https://github.com/nschloe/meshio

[70] M. König, S. Gerhäusser, SPT results - Simulation of zonation-function relationships in the liver using coupled multiscale models: Application to drug-induced liver injury (Apr. 2024). doi:10.5281/zenodo.10933166. URL 10.5281/zenodo.10933166

[71] D. Sasse, I. Maly, Studies on the periportal hepatotoxicity of allyl alcohol, Progress in Histochemistry and Cytochemistry 23 (1–4) (1991) 146–149. doi: 10.1016/s0079-6336(11)80180-4.

[72] S. A. Jung, Y.-H. Chung, N. H. Park, S. S. Lee, J. A. Kim, S. H. Yang, I. H. Song, Y. S. Lee, D. J. Suh, I.-H. Moon, Experimental model of hepatic fibrosis following repeated periportal necrosis induced by allylalcohol, Scandinavian Journal of Gastroenterology 35 (9) (2000) 969–975. doi: 10.1080/003655200750023057.

[73] E. E. Graham, R. J. Walsh, C. M. Hirst, J. L. Maggs, S. Martin, M. J. Wild, I. D. Wilson, J. R. Harding, J. G. Kenna, R. M. Peter, D. P. Williams, B. K. Park, Identification of the thiophene ring of methapyrilene as a novel bioactivation-dependent hepatic toxicophore, Journal of Pharmacology and Experimental Therapeutics 326 (2) (2008) 657–671. doi:10.1124/jpet.107.135483.

[74] K. B. Halpern, R. Shenhav, O. Matcovitch-Natan, B. Tóth, D. Lemze, M. Golan, E. E. Massasa, S. Baydatch, S. Landen, A. E. Moor, A. Brandis, A. Giladi, A. Stokar-Avihail, E. David, I. Amit, S. Itzkovitz, Single-cell spatial reconstruction reveals global division of labour in the mammalian liver, Nature 542 (7641) (2017) 352–356. doi:10.1038/nature21065.

[75] G. S. Ratra, W. A. Morgan, J. Mullervy, C. J. Powell, M. C. Wright, Methapyrilene hepatotoxicity is associated with oxidative stress, mitochondrial disfunction and is prevented by the Ca2+ channel blocker verapamil, Toxicology 130 (2–3) (1998) 79–93. doi:10.1016/s0300-483x(98)00096-1.

[76] G. S. Ratra, C. J. Powell, B. Park, J. L. Maggs, S. Cottrell, Methapyrilene hepatotoxicity is associated with increased hepatic glutathione, the formation of glucuronide conjugates, and enterohepatic recirculation, Chemico-Biological Interactions 129 (3) (2000) 279–295. doi:10.1016/s0009-2797(00)00253-2.

[77] Y. Yamaura, M. Nakajima, N. Tatsumi, S. Takagi, T. Fukami, K. Tsuneyama, T. Yokoi, Changes in the expression of mirnas at the pericentral and periportal regions of the rat liver in response to hepatocellular injury: Comparison with the changes in the expression of plasma mirnas, Toxicology 322 (2014) 89–98. doi:10.1016/j.tox.2014.05.008.

[78] M. Zhao, H. Xing, M. Chen, D. Dong, B. Wu, Circadian clock-controlled drug metabolism and transport, Xenobiotica 50 (5) (2019) 495–505. doi: 10.1080/00498254.2019.1672120.

[79] R. Craig Schnell, H. P. Bozigian, M. H. Davies, B. Alex Merrick, K. L. Johnson, Circadian rhythm in acetaminophen toxicity: Role of nonprotein sulfhydryls, Toxicology and Applied Pharmacology 71 (3) (1983) 353–361. doi:10.1016/0041-008x(83)90022-4.

[80] B. P. Johnson, J. A. Walisser, Y. Liu, A. L. Shen, E. L. McDearmon, S. M. Moran, B. E. McIntosh, A. L. Vollrath, A. C. Schook, J. S. Takahashi, C. A. Bradfield, Hepatocyte circadian clock controls acetaminophen bioactivation through nadph-cytochrome P450 oxidoreductase, Proceedings of the National Academy of Sciences 111 (52) (2014) 18757–18762. doi: 10.1073/pnas.1421708111.

[81] J. Grzegorzewski, J. Brandhorst, M. König, Physiologically based pharmacokinetic (PBPK) modeling of the role of CYP2D6 polymorphism for metabolic phenotyping with dextromethorphan, Frontiers in Pharmacology 13 (Oct. 2022). doi:10.3389/fphar.2022.1029073.

[82] H. Tautenhahn, T. Ricken, U. Dahmen, L. Mandl, L. Bütow, S. Gerhäusser, L. Lambers, X. Chen, E. Lehmann, O. Dirsch, M. König, SimLivA–modeling ischemia-reperfusion injury in the liver: A first step towards a clinical decision support tool, GAMM-Mitteilungen (Jan. 2024). doi:10.1002/gamm.202370003.

[83] M. Albadry, J. Küttner, J. Grzegorzewski, O. Dirsch, E. M. Kindler, R. Klopfleisch, V. Liska, V. Moulisová, S. Nickel, R. Palek, J. Rosendorf, S. Saalfeld, U. Settmacher, H.-M. Tautenhahn, M. König, U. Dahmen, Cross-species variability in lobular geometry and cytochrome P450 hepatic zonation: Insights into CYP1A2, CYP2E1, CYP2D6 and CYP3A4, bioRxiv (2023). URL https://api.semanticscholar.org/CorpusID:266694301

[84] A. Ghallab, M. Myllys, C. H. Holland, A. Zaza, W. Murad, R. Hassan, Y. A. Ahmed, T. Abbas, E. A. Abdelrahim, K. M. Schneider, M. Matz-Soja, J. Reinders, R. Gebhardt, M.-L. Berres, M. Hatting, D. Drasdo, J. Saez-Rodriguez, C. Trautwein, J. G. Hengstler, Influence of liver fibrosis on lobular zonation, Cells 8 (12) (2019) 1556. doi:10.3390/cells8121556.

[85] M. Albadry, S. Höpfl, N. Ehteshamzad, M. König, M. Böttcher, J. Neumann, A. Lupp, O. Dirsch, N. Radde, B. Christ, M. Christ, L. O. Schwen, H. Laue, R. Klopfleisch, U. Dahmen, Periportal steatosis in mice affects distinct parameters of pericentral drug metabolism, Scientific Reports 12 (1) (Dec. 2022). doi:10.1038/s41598-022-26483-6.

